# Exploring the culturable bacterial diversity and its hydrocarbon degrading potentiality isolated from the Oxygen Minimum Zone Sediments of Bay of Bengal

**DOI:** 10.1101/2022.06.05.494866

**Authors:** ChinnamaniKumar Prasannakumar

**Affiliations:** PG & Research Department of Biotechnology and Microbiology, National College (Autonomous), Dindigul road, Tiruchirappalli- 620 001, Tamil Nadu, India; Center of Advanced Study in Marine Biology, Faculty of Marine Science, Annamalai University, Parangipettai- 608502, Tamil Nadu, India

**Keywords:** Oxygen Minimum Zone, 16S rRNA gene, Bay of Bengal, sediment bacteria, hydrocarbonoclastics

## Abstract

Understanding biota distribution in oxygen minimum zone can help guide further exploration of potentially unusual habitats. The present study explores the culturable bacterial fractions in the oxygen minimum zone sediments of Bay of Bengal. The 16S rRNA gene sequences of 30 morphologically distinct bacterial colonies isolated form oxygen minimum zone of Bay of Bengal reveals 25 phylo-types, predominated by Proteobacteria (83.3%) and Actinobacteria (16.6%). Over all, Alphaproteobacteria and Gammaproteobacteria dominated the culturable fraction in this study. The overall pair-wise distances of bacterial isolates of Bay of Bengal is two times lesser when compared to overall pair-wise distance of bacterial isolates from oxygen minimum zone of Arabian Sea indicating relatively low genetic distances in Bay of Bengal. Not even 1% of bacterial cells in oxygen minimum zone of Bay of Bengal are culturable. We found that oxygen concentration alone could not be a deciding factor of culturable bacterial diversity in oxygen minimum zone. More than 50% bacterial isolates of present study is an active degraders of hydrocarbons. Higher similarity of 16S rRNA sequences produced in this study with that of previously reported efficient hydrobonoclastic bacterial isolates like *Vibrio diazotrophicus, Vibrio cyclotrophicus, Pseudomonas poae, Marinobacter hydrocarbonoclasticus, Marinobacter flavimaris* and *Alcanivorax borkumensis* further strengthens the evidence of hydrocarbon presence in Bay of Bengal sediments. This study is first of its time addresses the diversity of culturable bacterial fractions in oxygen minimum zone sediments of Bay of Bengal. Higher number of bacterial isolates from oxygen minimum zone of Bay of Bengal has carbonoclastic potentialities implying that they may play an important role in *in situ* hydrocarbon degradation in oxygen minimum zone of Bay of Bengal.

## 1. Introduction

Global Ocean may contain 1,148,000 km^2^ of Oxygen minimum zone, in which about 59% was contributed from Indian Ocean (Helly and Levin, 2004). Marine oxygen minimum zones (OMZ thereafter; <0.5 mL/L) encroach upon the continental margins at depths of 100–1,000 m along much of the Eastern Pacific Ocean, Arabian Sea and Bay of Bengal (BoB) (Levin 2003; Cowie and Levin 2009). They endure over geological time scales and result from a combination of factors, including high surface productivity and limited water-column ventilation caused by stratification and isolation of older, oxygen-depleted water masses. Organic matter concentrations, which typically are linked inversely to oxygen, are often high (Levin and Gage 1998) in OMZ. Hydrogen sulphide may be present within OMZ sediments but more often is removed by the formation of iron sulphides. The sediments (of OMZ) are often very soft and unconsolidated in the core regions of OMZ as a result of high water content. Variations in these parameters create gradients on the sea floor, rather than spatially distinct habitats. More discrete, visually obvious sources of heterogeneity are created by bacterial mats (Gallardo 1977; Schmaljohann et al. 2001) in OMZ. It is also possible that OMZ act as barriers that enhance diversity over evolutionary time by promoting genetic differentiation (Rogers 2000).

Bacterial mats often provide a localized, firmer, less toxic and geochemically distinct substratum within OMZs. On the Pakistan margin, a Beggiatoa / Thioploca mat yielded two foraminiferan species not present in adjacent sediments, as well as higher abundances of three other species and slightly higher diversity values compared to a core taken outside the mat (Erbacher and Nelskamp 2006). However, the overall abundances of stained benthic Foraminifera were much lower within the mat. Usually diversity among all taxa in OMZ is very low (Levin et al., 2001). Massive mats of sulfide-oxidizing bacteria may cover the sediments (Jorgensen and Gallardo, 1999) where symbiont-bearing invertebrates could occur (Cary et al. 1989; Levin et. al., 2002). Many OMZ species are new to science and some have exhibited unusual evolutionary novelty. For example, an inconspicuous gutless oligochaete discovered in the Peru OMZ, supports more types of nutritional bacterial symbionts (3 for certain and possibly 5) than reported from any other invertebrate (Giere and Krieger, 2001). There is growing evidence that chemosynthesis may fuel secondary production in OMZ transition zones where sulfide oxidation can occur (Levin 2003). An understanding of OMZ distributions can help guide further exploration of potentially unusual habitats. Though occupying larger proportions of global OMZ area, the biotic composition in OMZ of BoB is poorly studied (Helly and Levin, 2004). Hence the present study attempts to access the culturable bacterial fractions in the OMZ sediments of BoB.

## 2. Material and methods

### 2.1. Sample collection and bacteria isolation

Samples were collected during cruise no. 275 (May 2010) in research vessel Sagar Sampada. Samples were collected in two stations T1 and T4 along Eastern continental margins at the depths around 200m in BoB (fig. 1). Dissolved oxygen concentrations, temperature and pressure of water mass just above the sampled sediments were measured using Conductivity Temperature Depth profiler (CTD, SeaBird U.S.A.) available onboard. Collected sediments were immediately (not more than 4minutes after collection) subjected to dilution using polycarbonate Nuclepore membrane filter (GeNei, India) sterilized sea water and plated in Zobell Marine Agar 2216 (HiMedia, India). The plates incubated at 27_LJ_C were monitored at 24 hour interval for a period of 1 week in order to include isolation of the slow growing bacteria (Divya et al., 2010). Morphologically different isolates of T1 (n=16) and T4 (n=14) were sorted and purified for molecular and degradation studies.

**Fig. 1.**
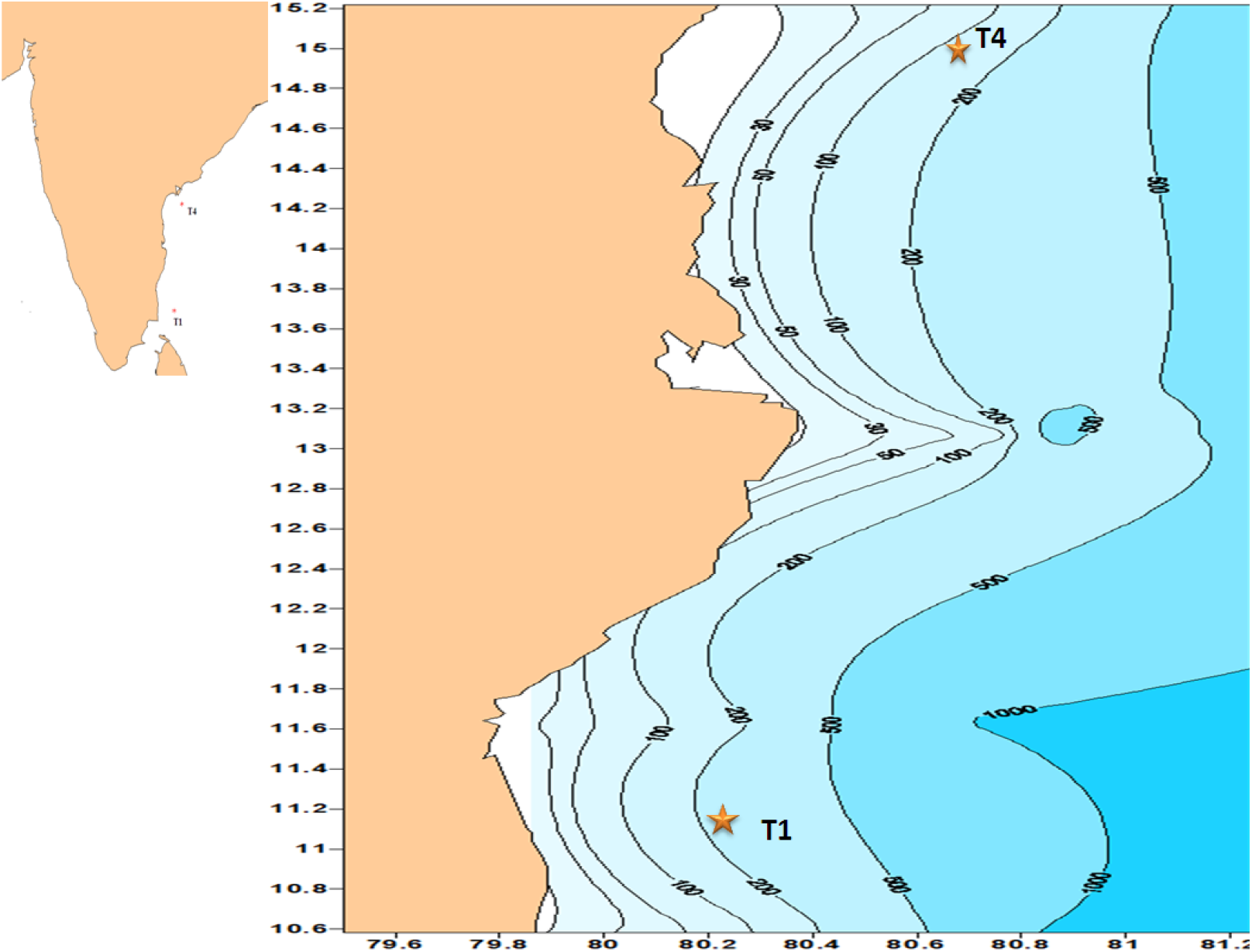
Map showing the sampling stations (T1 & T4; indicated by star mark) along Bay of Bengal. Depths between 150 to 1000m are known for oxygen minimal waters.

### 2.2. Total bacterial cell density

Total number of bacterial cells inhabiting sediments at the given time was estimated by Acridine Orange Direct count (AODC) (Hobbie et al., 1977). The sediments were sonicated for optimum removal of bacterial cells cling with sediments (Velji and Albright, 1986). The slides were prepared as soon as after sonication and always within 3hours of collection. Thirty random fields per slide were counted in an Olympus BHA epi-fluorescence microscope coupled with an image analysis system. The percentage of retrievability of the given transects could be calculated by dividing total numbers of bacterial counts (CFU) in general agar media divided by total number of bacterial cells counted via epi-fluorescence microscope, multiplied by 100 (for percentage conversion).

### 2.3. Polymerase chain reaction and 16S rRNA gene sequencing

For DNA extraction, to 50μL of double distilled water, a loop full of bacterial culture was dispensed and incubated in dry bath SLM-DB-120 (GeNei, India) maintained at 62□C for 30 minutes or loop full of culture in 50μL double distilled water vigorously vortexed (10x speed) and incubated at -20□C overnight. Following incubations, 1μL of slurry was used as template for polymerase Chain Reaction (PCR) and the primer pairs 8F (5’- AGAGTTTGATCATGGCTCAG-3’) and 1492R (5’-CGGTTACCTTGTTACGACTT-3’) (Lane, 1991) was used for 16s rRNA gene amplification, using thermal cycler model TC-3000 (GeNei, India). The PCR products were purified and sequenced through ABI 3770 at a commercial company, Bioserve Biotechnologies, India.

### 2.4. Organic carbons and hydrocarbon

For determining total organic content (TOC) of sediments, chromic acid oxidation method modified by Gaudette *et al*. (1974) was adopted. Total organic matter (TOM) contents were calculated using a conversion factor of 1.82 from TOC values. Petroleum hydrocarbon content of the sediments is estimated using American Public Health Association’s (APHA, 1989) protocol. Briefly, 1g of sediment was treated with hexane and optically analyzed using Varian make Cary Eclipse Spectrofluorometer (Agilent tech., U.S.A) at 310nm excitation and at 364nm emission. All the estimations were conducted in triplicates.

### 2.5. Hydrocarbon degrading potentialities

Crude oil was used as a source of hydrocarbons for testing the potentialities of OMZ isolates in hydrocarbon degradation (Pirnik et al., 1974). Briefly 10^8^ mL^-1^ of bacterial cells (from Zobell marine broth) were inoculated into mineral salt broth supplemented with 1% of crude oil as sole source of carbon. Gravimetric analysis was used for estimating crude oil degradation (Sakalle and Rajkumar, 2009). Briefly residual crude oil was extracted in preweighed flask with hexane in a separating funnel. Following extraction the residues were desiccated and weighed. Degradation ability of the strain was confirmed by the evidence of differences in weight of crude oil added before inoculation and recovered after incubation (Oloke and Glick, 2005).

### 2.6. Dry lab methodologies

The sequences were compared with other sequences in National Centre for Biotechnology Information (NCBI) database through Basic Local Alignment Searching Tool (BLAST) analysis [http://blast.ncbi.nlm.nih.gov/Blast] (Zhang et al., 2000). The 16S rRNA gene similarity (visible in BLAST search) equal to or more than 98% was considered to be same species (Stackebrandt and Goebel, 1994, Keswani and Whitman, 2001). The forward and reverse strands of sequenced DNA chromatograms were compiled and aligned using BioEdit (Hall, 1999). All compiled full length 16s rRNA gene sequences were multiple aligned in Clustal X ver. 2.0 (Larkin 2007). The 16S rRNA phylogram was constructed in Molecular Evolutionary Genetic Analysis (MEGA ver. 4.0) software (Tamura et al., 2007) using Kimura-2-parametric distance (Kimura, 1980) model. Pair-wise distance data was generated in MEGA using Kimura 2-parametric formulae and converted to percentage value (multiplied with 100) for better understanding.

### 2.7. DNA sequences

16S rRNA gene sequences (n=30) was submitted in GenBank and could be accessed through accession numbers, JQ045820-JQ045835, JQ045786-JQ045791, JQ068798-JQ073806.

## 3. Results

### 3.1. Distribution of bacterial phyla

Biotic composition of OMZ of Indian Ocean was poorly described and the OMZ of BoB has been rarely explored. Temperature, pressure and dissolved oxygen of OMZ range from 15 to 16.4□C, 193 to 198bar and 0.07 to 0.1mL/L, respectively. The previous studies on Arabian Sea (AS) OMZ exposed 28phylo groups of bacterial group dominated by Actinobacterial (AB) phyla whereas the present study reveals 25phylo groups dominated by Proeobacterial (PB) phyla in OMZ of BoB. However the phylogram reveals that bacterial lineages of AS OMZ is different from BoB, as sequences of AS claded away from 16S rRNA sequences produced from BoB (fig. 2). The 16S rRNA gene of 30 bacterial colonies of this study belonged to 25 phylo groups, predominated by PB (83.3%) and rest is contributed by AB (16.6%). Among proteobacterial lineages, Alpha- (APB) and Gammaproteobacteria (GPB) contributed 44% each, whereas Beta- (BPB) and Zetaproteobacteria (ZPB) contributed 4% and 8% respectively (fig. 3).

**Fig. 2.**
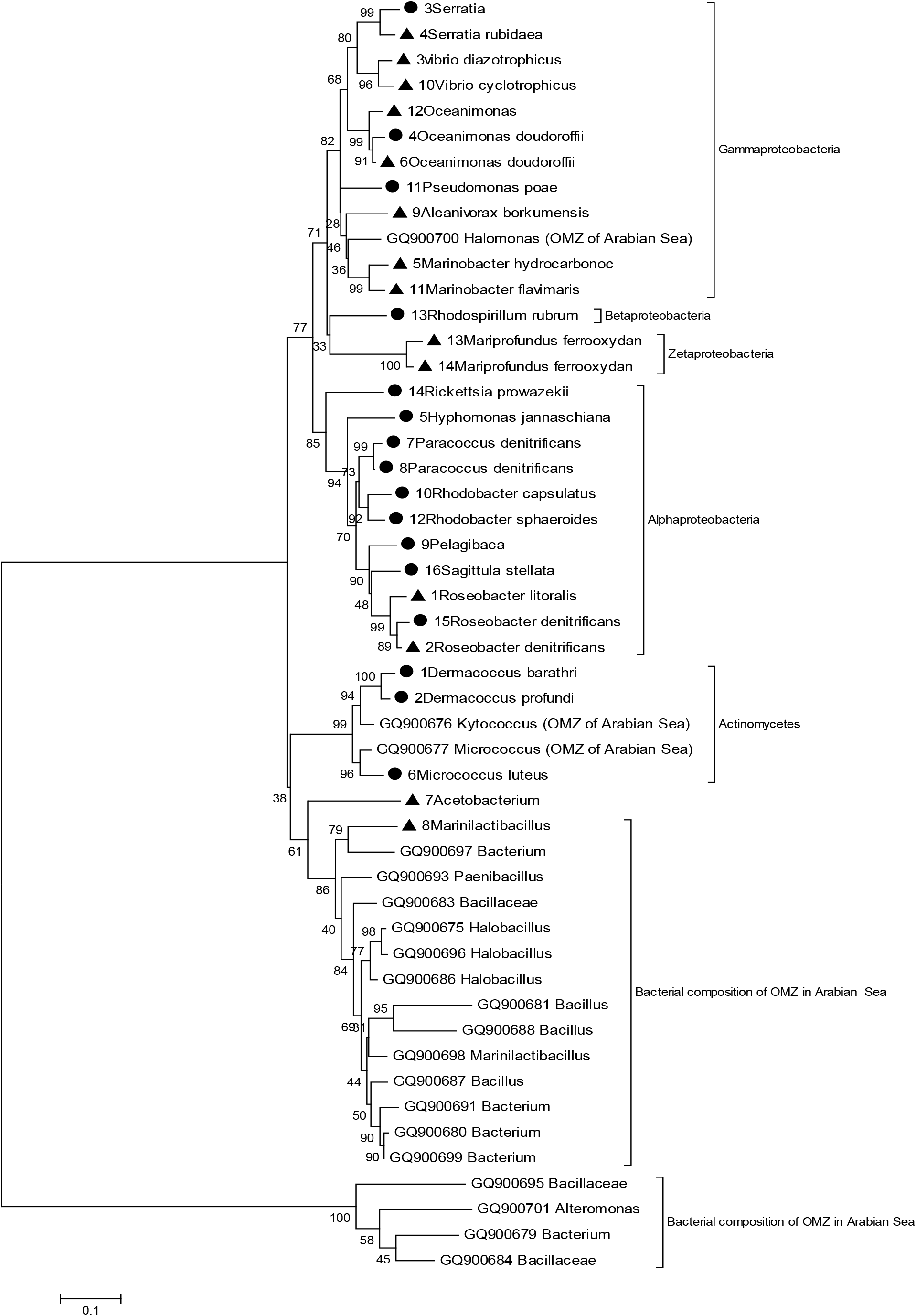
The evolutionary history was inferred using the Neighbor-Joining method. The optimal tree with the sum of branch length = 4.41297705 is shown. The percentage of replicate trees in which the associated taxa clustered together in the bootstrap test (500 replicates) is shown next to the branches. The tree is drawn to scale, with branch lengths in the same units as those of the evolutionary distances used to infer the phylogenetic tree. The evolutionary distances were computed using the Maximum Composite Likelihood method and are in the units of the number of base substitutions per site. All positions containing gaps and missing data were eliminated from the dataset (Complete deletion option). There were a total of 441 positions in the final dataset. Dark circles represents the identity of strains isolated in T1 transects, dark triangles represent strains isolated from T4 transect and the accession number before strain name indicate isolates form OMZ of Arabian Sea (Divya et al., 2010). Shorter 16S rRNA gene sequences (<500bps) in previous study (n=5) (Divya et al., 2010) was excluded from the analysis.

**Fig. 3.**
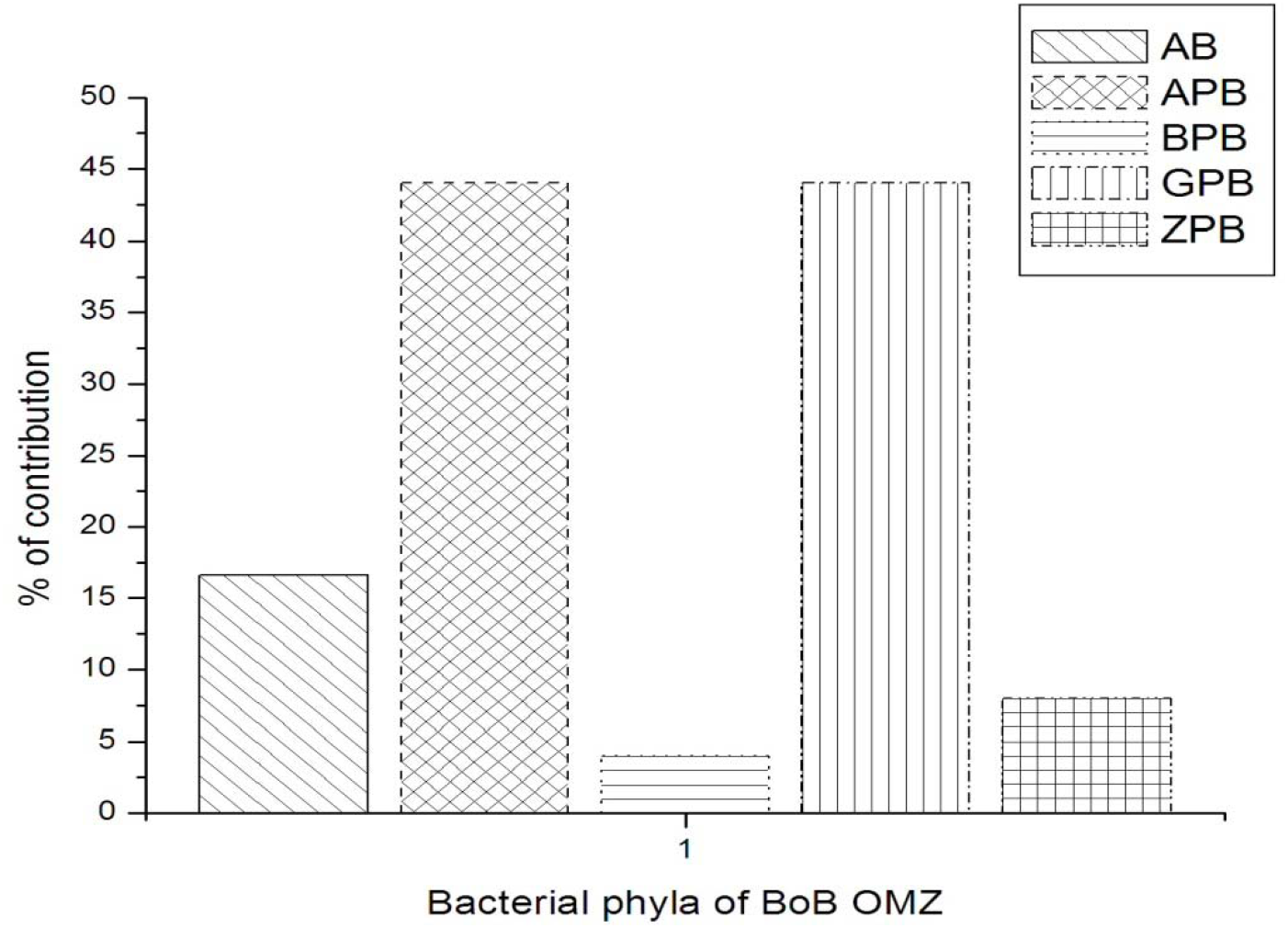
Graph representing percentage of contribution of various phylo-groups in culturable fractions of OMZ sediments of Bay of Bengal. AB-Actinobacteria, APB-Alphaproteobacteria, BPB-Betaproteobacteria, GPB-Gammaproteobacteria and ZPB-Zetaproteobacteria.

### 3.2. 16S rRNA gene diversity

We included all sequences (n=25) produced from OMZ of Arabian Sea for pair-wise distance analysis, along with 30 rRNA sequences produced in this study. The pair-wise distances between the isolates of T4 (0.252±0.015) is slightly higher than the isolates of T1 (0.248±0.014) transects. The overall pair-wise distances of bacterial isolates of BoB is found to be 0.254±0.013 which is still two times lesser when compared to overall pair-wise distance of isolates from OMZ of AS (0.594±0.042).

### 3.3. Phylogeny

Phylogram reveals that culturable bacterial fractions of OMZ in AS are completely different from that of BoB. Shorter 16S rRNA gene sequences (<500 base pairs (bps)) in previous study (n=5) was excluded from phylogenetic analysis. Among Arabian Sea isolates, AB group forms the major component followed by Firmicutes. But in PB group, especially APB and GPB dominated the culturable fraction of BoB OMZ sediments. Interestingly no Firmicutes (commonly found in Ocean waters) was recorded in this study. Phylogram contained two major clades. The PB classes formed a grouped in topmost clade (fig. 2) of the constructed phylogram. Other clade of phylogram contained AB with majority of isolates from OMZ of AS. Few members (n=4) of Bacillaceae family previously isolated from AS formed an out-group. The affiliation of isolates of AS to lower taxonomic level was difficult since most of the sequences produced in previous study was below 700bps (minimum of 273bp). Such difficulties were avoided in the present study as all 16S rRNA gene sequences (n=30) were of more than 1400bps length. Members of ZPB recovered from BoB is interesting one and its close relationship with members of BPB is reveled through cladding patterns of the phylogram (fig. 2). Among PB clades, APB was grouped separately from rest of the PB lineages recovered in this study.

### 3.4. Total bacterial cell count and retrievability

OMZ of BoB sediments were rich in bacterial density in the order of 10^8^cells/gram of sediments. Density of bacterial cells in T4 transects is 4 times higher than T1 transect (fig. 4). Likewise higher retrievability of bacterial colonies was witnessed from sediments of T4 (6.9×10^4^ CFU/g) transects than T1 (3.9×10^3^ CFU/g) transect which is more than 20 timer higher. But over all, not even 1% (by comparing total bacterial cells present and CFU recovered in ZMA) was of bacterial in OMZ of BoB is found culturable.

**Fig. 4.**
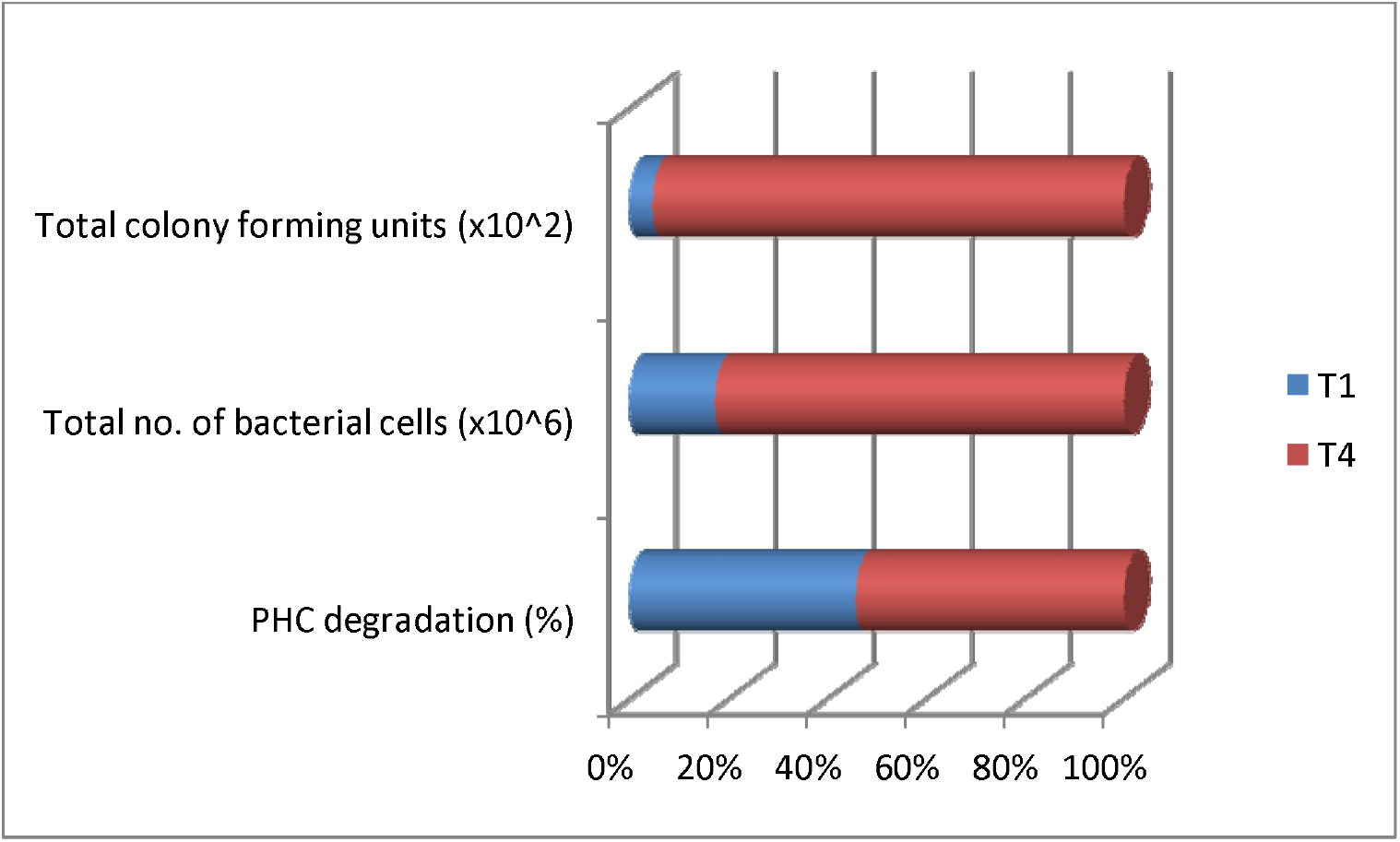
Percentage of bacterial cells found in sediments, percentage of cells recovered and percentage of CFUs exhibiting hydrocarbon degrading activity.

### 3.5. Hydrocarbon concentration and degradation

Total organic matter constitutes 5% of sediment weight. Transect T4 contained relatively higher amount of petroleum hydrocarbons than T1 (fig. 4). Over all 2.27 μg/g of petroleum hydrocarbons occurs in the sediments of OMZ. Interestingly more than 50% of isolates recovered in this study possessed hydrocarbon degrading potentialities. Transect T4 contained 62.8% and T1 contained 53% of hydrocarbon degrading bacterial phyla. Well known hydrocarbon degrading species like *Marinobacter hydrocarbonoclasticus* and *M. flavimaris* was retrieved in T4 transect. T1 transect contained *Pseudomonas poae* as a known hydrocarbon degrading candidate.

## 4. Discussion

Global warming may lead to lowered oxygen content of the world oceans (Keeling and Garcia, 2002), and expansion of OMZs in selected areas. Because OMZs are inhospitable to many species, they serve as biogeographic barriers, limiting cross-slope movements of populations (White, 1987; Etter et al., 1999; Rogers, 2000; Weeks et al., 2002). OMZ expansion or shrinkage may promote the evolution of species and genetic diversity maxima at mid-slope depths (Jacobs and Lindberg, 1998; Etter et al., 1999; Ulloa et al., 2001). The extent and severity of OMZs will change with alteration of ocean circulation, temperature and productivity (Reichart et al., 1998; Keeling and Garcia, 2002). Documenting the nature of microbial flora in such environment would facilitate predication of changes in these environments, there by easies better management protocols. This study for first of its time address the diversity of culturable bacterial flora in two transects along OMZ sediments in Indian part of Bay of Bengal.

### 4.1. Bacterial phyla distribution in OMZ

Notwithstanding the prospective of cultivation-independent techniques to provide taxonomic identification; in this study we have adopted pure culture technique to link bacterial species with their hydrocarbon degrading functions. In the present study, 30 sequences of 16S rRNA gene were generated belonging to 25 phylogroups of culturable bacterial isolates from sediments of OMZ of BoB. Predominance of PB was witnessed followed by AB in the OMZ sediments. PB constitutes only 50% among the culturable isolates of AS (Divya et al., 2010) whereas 83.3% of BoB OMZ sediments were occupied by PB lineages. Among PB group, APB and GPB dominated OMZ sediments which was also the case in the deep sea sediments of Pacific Ocean (Kato et al. 1996; DeLong et al. 1997; Wang et al. 2004). Interestingly all sequences in the present study could be classified to lower taxonomic levels whereas more than 62% of OMZ isolates of AS could not be classified (Divya et al., 2010). However few sequences showed lesser similarity to all known sequences in the database and whence the possibility of novel species endemic to this region could not be ruled out.

Though earlier study in Pacific Ocean has reported low species richness (Levin 2003), cultivation-independent studies of the sediments in OMZs of Pacific and Atlantic oceans have shown enormous bacterial diversity (Schippers and Neretin 2006; Liu et al. 2003 a, b). Further, this technique has facilitated the detection of uncultured bacteria which at times constitute 99% of the total bacteria (Olsen 1986; Schloss and Handelsman 2004). This was reiterated in the present study, as not even 1% of bacterial cells in OMZ of BoB were found culturable. Though bacterial cells of high orders (10^8^cells/g) were witnessed through AODC methods, maximum of 10^4^ CFU/g were witnessed in plate count technique reiterating “great plate count anomaly” (Staley and Konopka, 1985).

Alteration of ocean circulation, temperature and productivity (Keeling and Garcia, 2002) along with expansion or shrinkage (Etter et al., 1999) will change the extent and severity of OMZ that will promote the evolution of species and genetic diversity maxima in these regions (Ulloa et al., 2001). The pair-wise distance analysis of 16S rRNA gene revealed that culturable isolates in OMZ are more diverse than those in BoB. The pair-wise distance of 16S rRNA gene sequences of AS OMZ (Divya et al., 2010) were 2 times higher when compared with 16s rRNA gene sequence in the present study. However sequences of previous study were of shorter (all were <1000bps) length and whence defined conclusion on genetic distances were not probable.

*Marinobacter* and *Pseudomonas* were dominant genus among the culturable isolates of BoB OMZ, where *Bacillus* dominates the OMZ of Arabian Sea (Divya et al., 2010). It is becoming clear that oxygen concentration alone could not be a deciding factor of culturable bacterial diversity in a given area, as the OMZ sediment of Okinawa Trough and southwestern Okhotsk Sea where dominated by *Halomonas* spp. (Dang et al. 2009; Inagaki et al. 2003). Occurrences of a few dominant phylotypes reported form different areas of Ocean (Giuliano et al. 1999; Uphoff et al. 2001) may be due to habitat adaptations enabling them to survive in the OMZ environment. The total bacterial counts of U.S.A. waters vary from 10^5^ to 10^6^cells/mL (Rappe et al., 2002) where similar patterns are witnessed in OMZ waters of Arabian Sea (Goltekar et al., 2006), but the sediments from BoB OMZ was found to harbor high concentrations of bacterial cells which was in the order of 10^8^cells/g. This may be due to nature of sediment texture of this environment as the distribution of benthic communities is highly correlated with the type of sediment, which is related to a wider set of environmental conditions, such as current speed and organic content of the sediment (Buchanan 1984). Thus benthic assemblages are associated with grain size of the sediments. Silty nature of sediments were common in AS (Reddy, 2003), where as BoB shelf region are sandy in nature (Ganesh and Raman 2007) which may have profound influence on the bacterial concentrations.

### 4.2. Hydrocarbon degradation and ecological importance

The significance of metabolically active aerobic culturable bacteria of OMZ in cycling of organic matter has been reiterated in previous study (Divya et al., 2010). The OMZ sediments of the AS are found to harbor diverse bacteria which have enormous catabolic efficiencies. The supply of organic material to the OMZ sediment is a key factor in stimulating the production of extracellular hydrolytic enzymes (Divya et al., 2010). These hydrolytic enzymes are responsible for organic matter mineralization and thereby play important role *in situ* biogeochemical processes. BoB is considered organically poor when compared to AS (Khan et al., 2011), but was richer in hydrocarbon deposits (Paropkari et al., 1992, Lyla et al., 2012). We found that more than 50% bacterial isolates of OMZ sediments in BoB is an active degraders of hydrocarbons. T4 transects had higher number (62%) hydrocarbonoclastic bacterial isolates than T1 transect. The abundance of hydrocarbon degrading bacteria is may be due to natural occurrence of hydrocarbon deposits in the sediments of Bay of Bengal (Paropkari, 2008, Lyla et al., 2012). The TOC levels recorded in this study is concordant with the previous analysis (Khan et al., 2011). Higher similarity of 16S rRNA sequences produced in this study with that of previously reported efficient hydrobonoclastic bacterial isolates like *Vibrio diazotrophicus* (Guerino et al., 1982), *Vibrio cyclotrophicus* (Hedlund and Staley, 2001), *Pseudomonas poae* (Behrendt et al., 2003), *Marinobacter hydrocarbonoclasticus* (Gauthier et al., 1992), *Marinobacter flavimaris* (Yoon et al., 2004), *Alcanivorax borkumensis* (Kasai et al., 2002) further strengthens the evidence.

## 5. Conclusion

The significance of metabolically active aerobic culturable bacteria in cycling of hydrocarbon was reiterated in this study. The OMZ sediments of the BoB harbor diverse bacterial phyla which were found to have enormous hydrocarbon catabolic efficiencies. However the sources of hydrocarbon (anthropogenic or abiogenesis) in sediments are still a debatable subject in case of BoB. Higher number of bacterial isolates from OMZ of BoB has carbonoclastic potentialities implying that they may play an important role in *in situ* hydrocarbon degradation in OMZ of BoB.

---

**Table 1:**
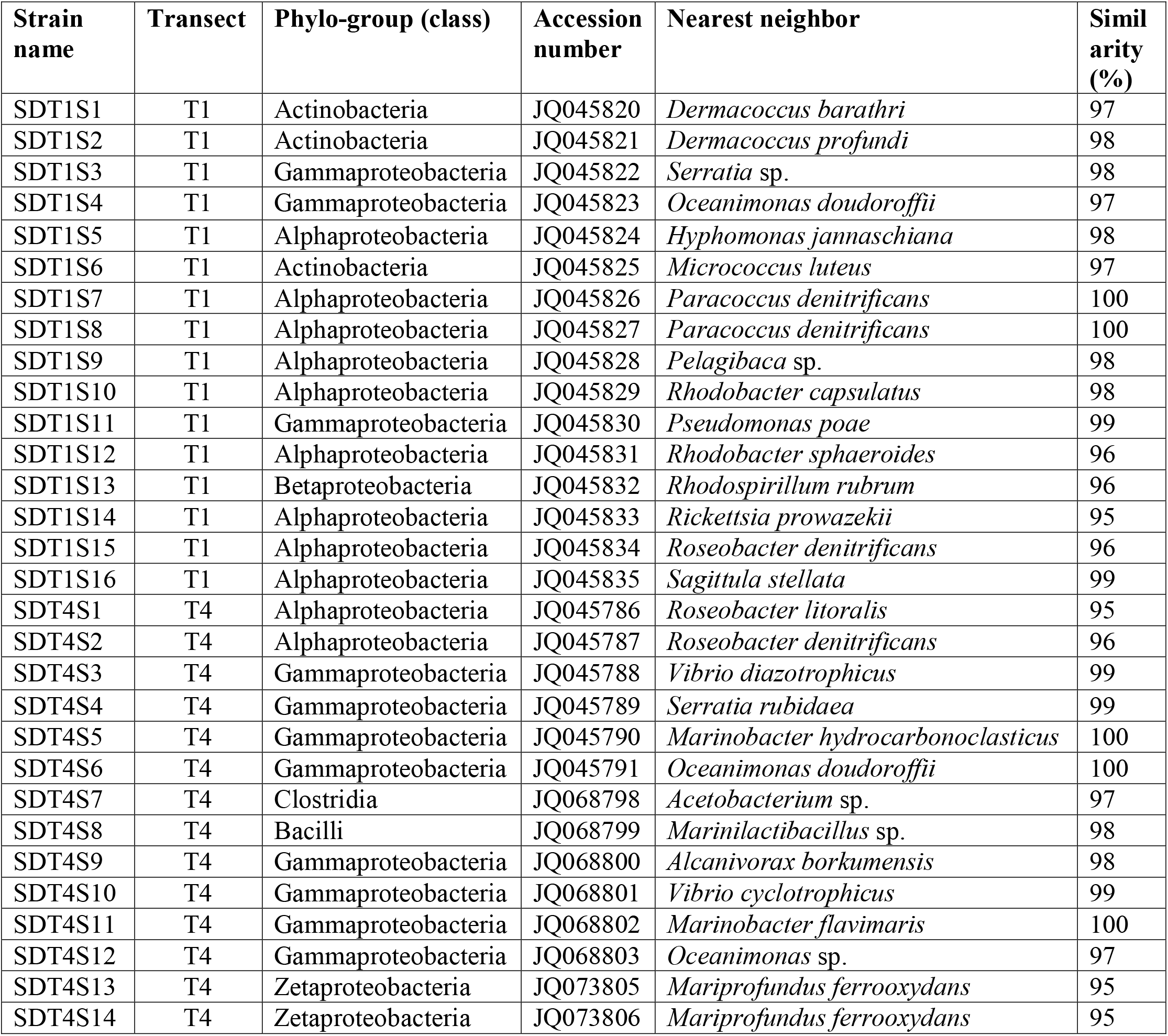
Nearest neighbors of isolated strain in GenBank

**Table 2:** No. of phylo-groups shared between transects

|  | T1 | T4 |
| --- | --- | --- |
| No. of 16S rRNA sequenced | 16 | 14 |
| No. of Phylogroups | 15 | 13 |
| Pair-wise distances | 0.248±0.014 | 0.252±0.015 |
| Total organic matter (%) | 4.5 | 5.4 |
| PHC concentration in sediments (μg/g) | 1.81 | 2.72 |
| PHC degradation (%) | 53 | 62.8 |
| Total no. of bacterial cells (cells/g) | 2.03$\times 10^8$ | 9.69 $\times 10^8$ |
| Total colony forming units (CFU/g) | 3.4$\times 10^3$ | 6.9$\times 10^4$ |

